# A new strategy for identifying mechanisms of drug-drug interaction using transcriptome analysis: Compound Kushen injection as a proof of principle

**DOI:** 10.1101/592956

**Authors:** Hanyuan Shen, Zhipeng Qu, Yuka Harata-Lee, Jian Cui, Thazin Nwe Aung, Wei Wang, R. Daniel Kortschak, David L. Adelson

## Abstract

Drug-drug interactions (DDIs), especially with herbal medicines, are complex, making it difficult to identify potential molecular mechanisms and targets. We introduce a workflow to carry out DDI research using transcriptome analysis and interactions of a complex herbal mixture, Compound Kushen Injection (CKI), with cancer chemotherapy drugs, as a proof of principle. Using CKI combined with doxorubicin or 5-Fu on cancer cells as a model, we found that CKI enhanced the cytotoxic effects of doxorubicin on A431 cells while protecting MDA-MB-231 cells treated with 5-Fu. We generated and analysed transcriptome data from cells treated with single treatments or combined treatments and our analysis showed that opposite directions of regulation for pathways related to DNA synthesis and metabolism appeared to be the main reason for different effects of CKI when used in combination with chemotherapy drugs. We also found that pathways related to organic biosynthetic and metabolic processes might be potential targets for CKI when interacting with doxorubicin and 5-Fu. Through co-expression analysis correlated with phenotype results, we selected the MYD88 gene as a candidate major regulator for validation as a proof of concept for our approach. Inhibition of MYD88 reduced antagonistic cytotoxic effects between CKI and 5-Fu, indicating that MYD88 is an important gene in the DDI mechanism between CKI and chemotherapy drugs. These findings demonstrate that our pipeline is effective for the application of transcriptome analysis to the study of DDIs in order to identify candidate mechanisms and potential targets.

## Introduction

Drug combinations or polypharmacy is a commonly used clinical strategy for elderly patients and chronic diseases like diabetes, cardiovascular disease and cancer, in order to overcome unwanted off-target effects and compensatory mechanisms for certain drugs ^123^. However, the challenge for polypharmacy ^4^ is how to estimate the effects of drug combinations compared to single drugs, and avoid potentially serious adverse effects resulting from drug-drug interactions (DDIs). The most common strategy for identifying DDIs is through pharmacokinetic approaches. This is because, by affecting transporters and metabolizing enzymes, one drug’s pharmacokinetic process (absorption, distribution, metabolism or excretion) can be changed by another drug. However, pharmacokinetic properties are not usually directly linked to pharmacodynamic effects and cannot show interactions with treatment targets or potential side effects. Furthermore, pharmacodynamic assays may not provide enough information for detecting potential interaction effects and interpreting their mechanisms ^5^. This is a particular concern for drug interactions involving complementary and alternative medicines (CAM), where herbal extracts can contain over a hundred different, potentially bioactive, compounds.

Public acceptance of combining complementary and alternative medicine (CAM) with conventional medicines has increased significantly over the last few decades. In 2007, nearly 38% of American adults used CAM ^6^, and in China, which has a long history of traditional herbal medicine, 93.4% of cancer patients use CAM ^7^. These medicines, especially traditional Chinese medicines (TCM) which are usually made from several herbs, can also exert their effects on conventional medicines both through pharmacokinetic and pharmacodynamic effects. The complexity of components in these CAMs make it extremely difficult to predict potential interactions with conventional medicines and explain these rationaly. By providing opportunities to examine a broad range of biological information, omics-related techniques provide a more comprehensive way for the study of drug-drug or herb-drug interactions ^8^. In this report, we apply these methods to the identification of interactions between Compound Kushen Injection (CKI), a complex herbal extract mixture, and chemotherapy drugs.

In this study CKI is used as a model complementary medicine. CKI was approved by the State Food and Drug Administration (SFDA) of China in 1995, CKI is used by more than 30,000 patients every day as part of their treatment for various types of cancers ^9^. Previous reports have shown that CKI can sensitize cancer to chemotherapeutic drugs, and reduce side effects of chemotherapy and radiotherapy to improve treatment effects and quality of life for cancer patients ^10,11^. CKI is extracted from two herbs, Kushen (*Radix Sophorae flavescentis*) and Baituling (*Rhizoma Smilacis glabrae*), which contain many natural compounds including, but not limited to alkaloids and flavonoids. Matrine and oxymatrine have been implicated as the primary active components for cancer treatment ^12^, but this is not supported by our previous research that showed that CKI, but not oxymatrine, can inhibit cancer cell proliferation and cause apoptosis by perturbing the cell cycle and other cancer related pathways ^13–15^. However, to date, no reports have revealed how CKI or its active components interact with cancer chemotherapy drugs.

In order to better understand DDIs and deal with the difficulties caused by complex components in herb-drug interactions, we propose a pipeline to apply transcriptome analysis for the study of DDIs. CKI was used as a test drug in combination with different chemotherapy agents, and was found to have different effects on cancer cells when combined with doxorubicin or 5-Fu (synergistic with doxorubicin and antagonistic with 5-Fu). Based on transcriptome data, we have identified hundreds of differentially expressed genes that are correlated with opposite effects of CKI and chemotherapy agents on cell viability or apoptosis. These genes indicate that several cancer related pathways, such as DNA replication and cell cycle, are perturbed differently by CKI under different medical circumstances. Compared to previous DDI studies focused on transporters, metabolizing enzymes and therapy targets, our methods can provide a comprehensive and deeper analysis of interactions, that may help to pinpoint potential therapeutic or side effects, and explain the mechanisms underlying DDIs.

## Results

### Pipeline for the study of DDIs using transcriptome analysis

Figure 1 shows the flowchart for transcriptome analysis of DDIs. First, assays for DDIs are selected that are suitable for RNA sequencing and phenotype readouts. Second, shared DE genes from treatment with the primary drug only and combined treatment of primary drug and interacting drug are identified and further classified based on their manner of regulation. Gene co-expression analysis can then be used to identify groups of genes whose regulation is correlated with phenotype. Finally, different annotation methods can be used to propose mechanisms for DDIs and predict potential interactions.

**Figure 1:**
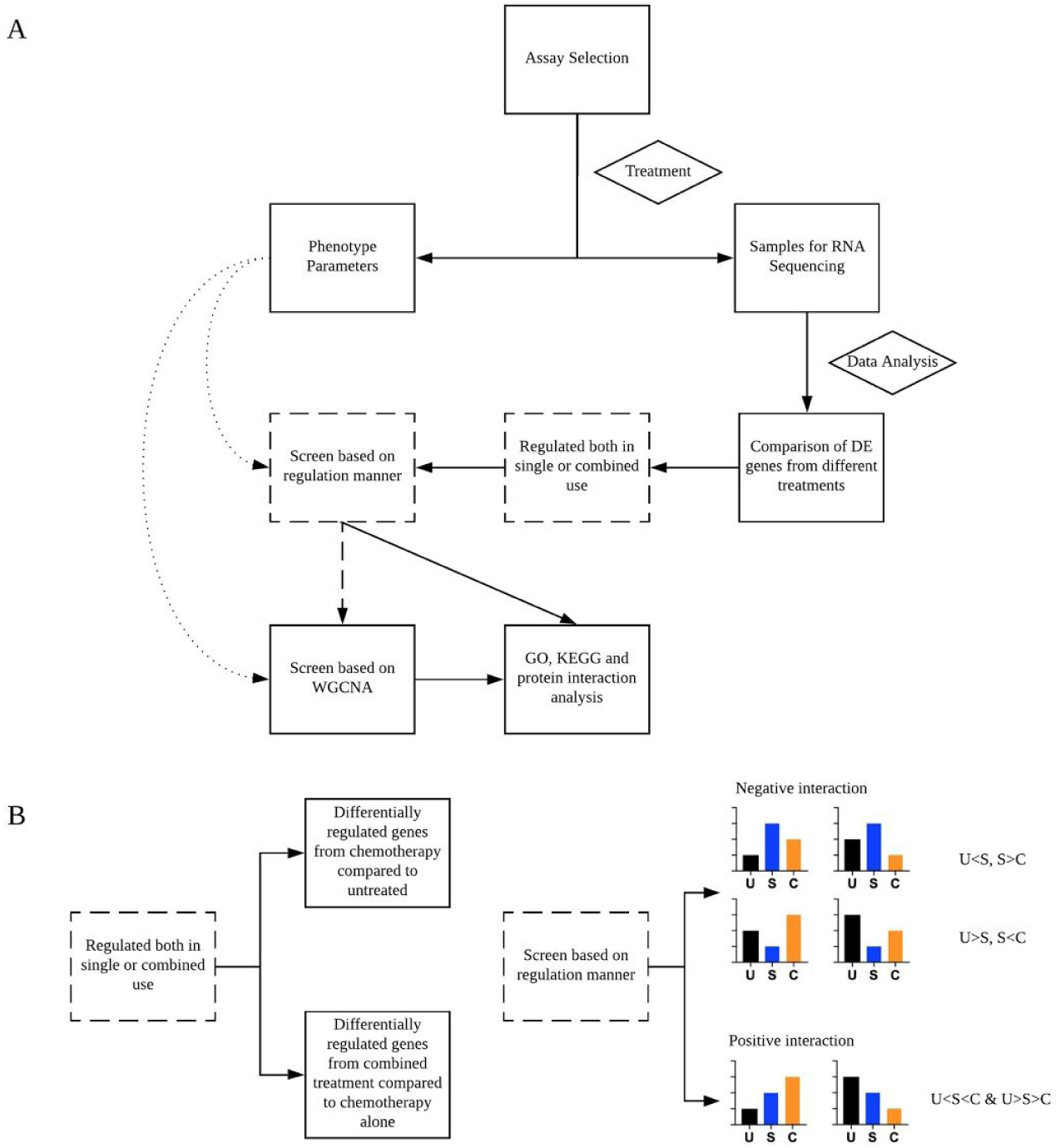
Experimental and data analysis workflow for applying omics to drug-drug interactions. A. The overall design of the study. B. Further details of 2 specific procedures indicated with broken-lined boxes in A. The black, blue and orange bars represent untreated, single treatment and combined treatment, respectively.

The differentially regulated genes for single drug treatment are calculated with respect to untreated samples, while combined treatments are compared to single treatment. In addition, to identify types of interactions, genes consistently up or down regulated in single treatment and in combined treatment are classified as positively interacting (in other words, the expression level of from primary chemotherapy agent treatment is intermediate between untreated and combined treatment). Negatively interacting genes have expression levels where the primary chemotherapy treatment causes either the highest or lowest expression compared to untreated cells, or combined treatment).

### CKI enhances the effects of doxorubicin but protects cells when co-administered with 5-Fu

CKI alone can inhibit proliferation, induce apoptosis and alter the cell cycle for various cancer cell lines ^13–15^. In order to determine whether CKI can potentiate the anticancer effects of chemotherapy agents, we used the XTT assay as a preliminary screen for the interaction of CKI with different chemotherapy drugs (Supplementary Fig. 1). Results showed that CKI could have opposite effects in different chemical contexts. These effects were most obvious at relatively low doses of CKI and chemotherapy agents to treat MDA-MB-231(with 5-Fu) and A431 cells (with doxorubicin) for 48 hours. CKI increased the apoptotic effects of doxorubicin whereas it antagonized the cytotoxicity of 5-Fu (Fig. 2A). Flow cytometric analysis of propidium iodide (PI) stained cells was also used to assess alterations to the cell cycle and apoptosis for different treatment groups. In MDA-MB-231 cells, treatment with CKI caused the increased percentages of apoptosis from 5-Fu to be drawn back to the same level as untreated cells. However, the proportion of apoptotic cells increased significantly when CKI was combined with doxorubicin on A431 cells (Fig. 2B). Also, compared to slight changes in the cell cycle caused by the combination of CKI and 5-Fu, CKI caused large decreases in G1 and S phases of the cell cycle compared to doxorubicin only treatment (Fig. 2C). Altogether, these data suggest that CKI has opposite interactions with doxorubicin and 5-Fu *in vitro*.

**Figure 2:**
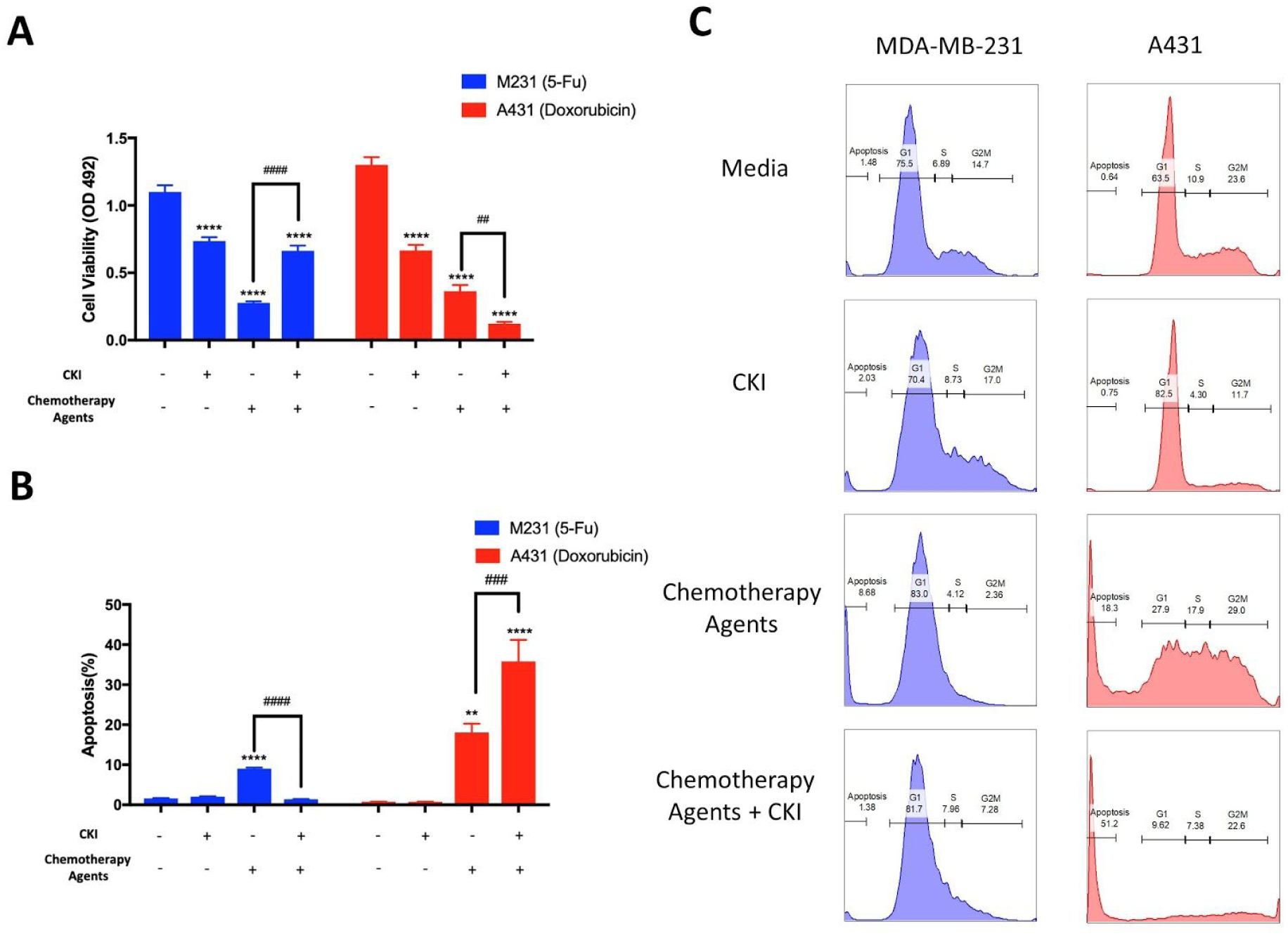
Opposite effects of CKI combined with doxorubicin or 5-Fu on cell viability and cell cycle. A. and B. The cell viability and percentage of apoptosis of the cancer cells treated with different drug combinations for 48 hours. C. Representative histograms of cell cycle phases for different treatments. Results are represented as means ±SEM (n=9). Statistical analyses were performed by comparing treatments to untreated (**p< 0.01, ***p < 0.001, **** p < 0.0001) as well as ‘CKI + Chemotherapy Agent’ to ‘Chemotherapy Agent only’ (##p<0.01, ###p<0.001, ####p<0.0001) with one-way ANOVA.

### Selecting DE genes involved in drug-drug interactions

In order to understand the molecular mechanisms of the opposite interactions of CKI with doxorubicin and 5-Fu, we carried out transcriptome profiling from chemotherapy agents treatment, CKI treatment and combined CKI+chemotherapy using high-depth next generation sequencing. In order to correlate the gene expression results with phenotype results, we selected 48 hours as the treatment time with three biological replicates. After preliminary multidimensional scaling of all the samples, every treatment group clustered, and clusters were clearly separated, indicating that combined CKI treatment can change the transcriptome of cancer cells compared to chemotherapy or CKI alone (Supplementary Fig. 3 & 4).

Because we were primarily interested in determining the changes in gene expression between combined and single treatments, we identified DE genes by comparing the combined treatment to treatment with chemotherapy drug only. We also compared single treatments to untreated. This gave 4 sets of DE genes (A431 cell line: doxorubicin compared to untreated and doxorubicin + CKI compared to doxorubicin, MDA-MB-231 cell line: 5-Fu compared to untreated and 5-Fu + CKI compared to 5-Fu) with each set containing thousands of DE genes (Fig. 3A & Supplementary Table 2).

**Figure 3:**
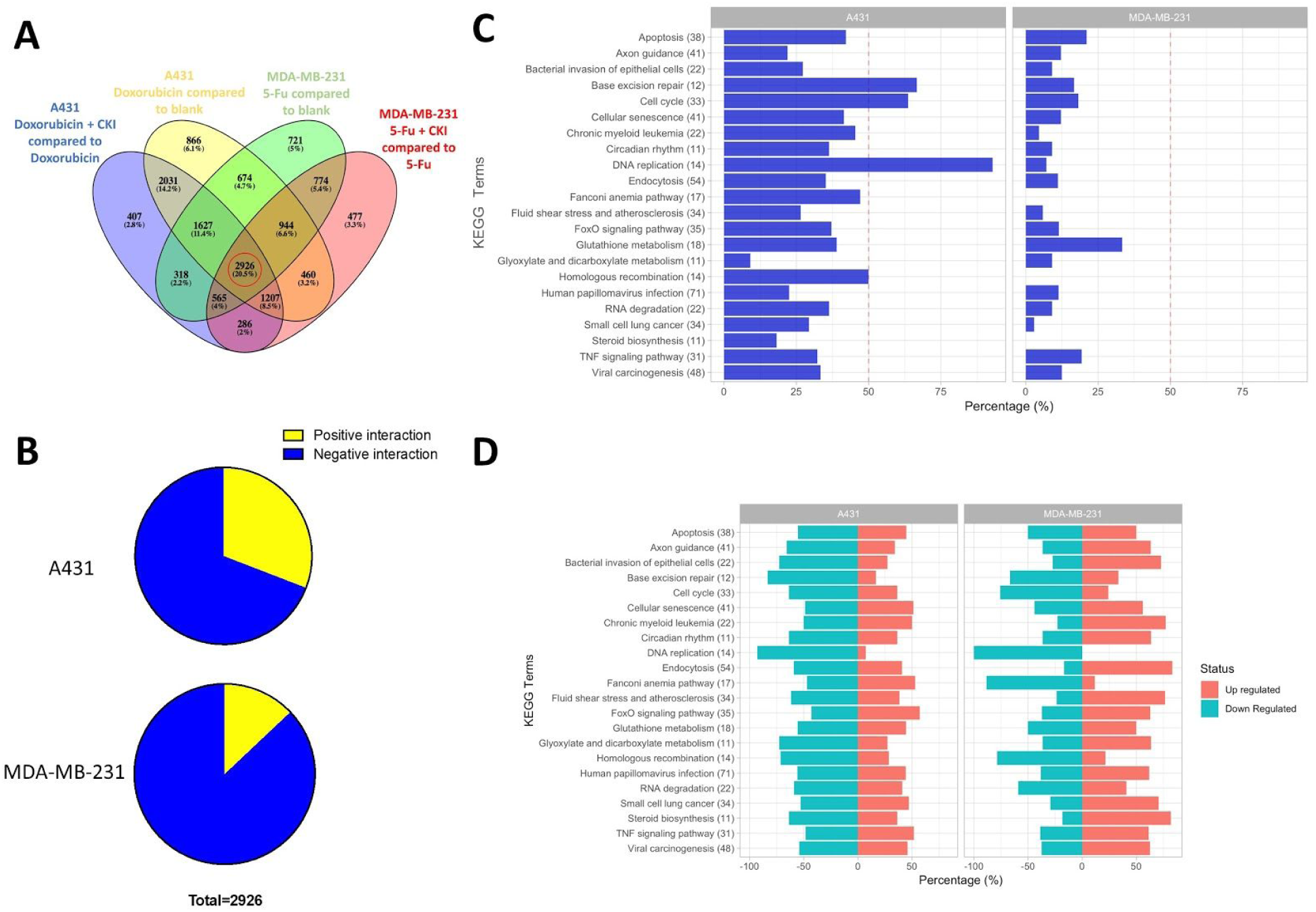
Selection of differentially regulated shared genes and percentage of genes regulated in different fashions and their related pathways. A. Venn diagram showing the number of differentially regulated genes in cancer cells with different treatments. B. Percentage of genes that were regulated in ‘synergistic’ (yellow) and ‘antagonistic’ (blue) fashion in A431 (doxorubicin) and MDA-MB-231 (5-Fu) cells. C. Percentage of synergistically regulated genes in different KEGG pathways. D. Percentage of up-regulated and down-regulated genes for single chemotherapy drug treatment in different KEGG pathways.

### Identification of DE genes based on interaction and direction of regulation

From the original four gene sets, we refined our results to get DE gene subsets related to drug interactions. Because of our primary focus on the mechanisms underlying opposite effects of CKI combined with doxorubicin or 5-Fu, we identified the set of common DE genes across the four sets of DE genes identified above. This subset of 2926 genes was selected for further analysis (Fig. 3A). Because differential expression can result from up or down regulation of expression, we included the direction of interaction as a means to separate DE genes involved in DDIs. If one gene’s change in expression level from untreated to single treatment was consistent with its direction of regulation for the combined treatment (either up regulated or down regulated) then we defined it as a positive interaction gene, either additive or synergistic. In contrast, if its direction of regulation was opposite in the single treatment compared to the combined treatment, it was defined as a negative interaction. When these criteria were applied to our subset of 2926 genes, while most of the DE genes underwent negative interaction across both cell lines/treatments, the proportion of positive interaction genes differed between treatment groups. In A431 cells treated with doxorubicin, CKI induced 30.9% positive interaction genes, whereas only 12.9% of the genes were positively interacted with CKI in MDA-MB-231 cells treated with 5-Fu (Fig. 3B).

In order to further characterise the genes with different directions of interaction, we performed functional enrichment analysis of Kyoto Encyclopedia of Genes and Genomes (KEGG) pathways for our set of shared 2926 DE genes and calculated the number of genes for negative and positive interaction with CKI in both treatment groups/cell lines. The results showed that in every pathway, the proportion of DE genes positively regulated by CKI treatment with doxorubicin (A431 cells) was larger than by CKI treatment with 5-Fu (MDA-MB-231 cells) (Fig. 3C). Strikingly, there were four pathways related to cell cycle that had over 50% the genes positively regulated in the A431 cells, including: “Base excision repair”, “Cell cycle”, “DNA replication” and “Homologous recombination”. When we took into account the expectation that one third of the DE genes should fall into the positive interaction class (Fig. 1B), we used 33.33% as the cut off for distinguishing direction of interaction pathways. With this criterion, there were 13 pathways where CKI caused positive interactions with doxorubicin, but only 1 with 5-Fu (Fig. 3C). Furthermore, 9 pathways were consistently found to interact in a positive manner, including immune pathways (“Bacterial invasion of epithelial cells”, “Human papillomavirus infection”, “Viral carcinogenesis”) and metabolic pathways (“Glyoxylate and dicarboxylate metabolism”, “Steroid biosynthesis”) and others..

To have a comprehensive understanding of drug-drug interactions, samples treated with single chemotherapy agents were also annotated with KEGG pathways. With the common regulated subset, genes in 8 cell cycle related pathways were primarily regulated in the same directions (mainly down-regulated in 6 pathways and up-regulated in 2 pathways) both by doxorubicin and 5-Fu (Fig. 3D).

### DE genes related to phenotype

Based on the direction of regulation in each combined treatment group, we separated the 2926 shared DE genes into four groups (Fig. 4A & Supplementary Table 3). Group C in which genes were negatively interacted in both cell lines, contained the largest number of genes (1815, 62% for total gene number) followed by group A genes with 732 that are negatively interacted for 5-Fu and positively interacted for doxorubicin. The other two groups (C and D) only contained 208 and 171 genes respectively. Based on the phenotype results, genes in group A were more likely to relate to our study purpose, while groups C and D might reveal CKI’s overall effects on chemotherapy cancer drugs.

**Figure 4:**
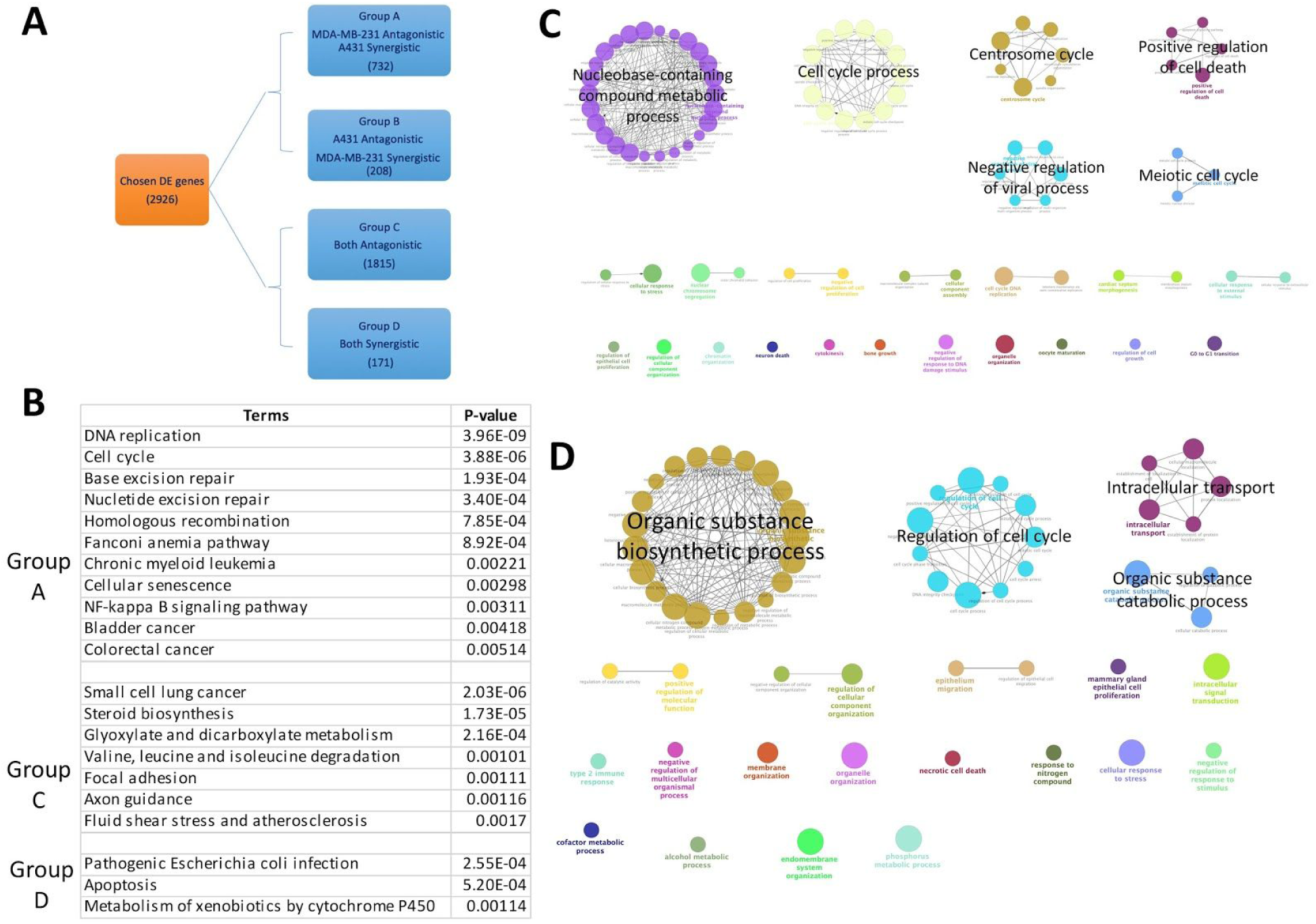
Grouping of genes based on type of regulation and annotation results for different gene groups. A. Criteria for separating 2926 genes into four groups. B. Table for over-represented KEGG terms and associated p-values for different groups of genes. C. and D. Over-represented GO terms (Biological Process at 3rd level) for genes in groups A and C.

Functional enrichment analysis was also performed for each of the four groups (Fig. 4B, C & D & Supplementary Table 3). For group A, the Gene Ontology terms for 732 genes (Fig. 4C) were mainly related to “cell cycle” and “nucleobase-containing compound metabolic process”. In KEGG pathway analysis, except for pathways closely related to cell growth like “Cell Cycle” and “DNA replication”, there were two cancer related pathways detected, “Bladder cancer” and “Chronic myeloid leukemia” and one immune related pathway “NF-kappa B signaling pathway”. Genes in these pathways, like cell cycle (Fig. 5A & Supplementary Fig. 2) are regulated in opposite direction by CKI compared to chemotherapy drugs. GO terms for 1815 genes in group C (Fig. 4D) were mainly clustered into “Organic substance biosynthetic process”, “regulation of cell cycle” and “organic substance catabolic process”, KEGG results also indicated that the majority of genes in this group belonged to pathways related to metabolism and biosynthesis. Gene numbers for groups B and D were much lower, with mainly immune related GO terms for group B and cell cycle related GO terms for group D. Only three KEGG pathways were significantly enriched for group D and none for group B.

**Figure 5:**
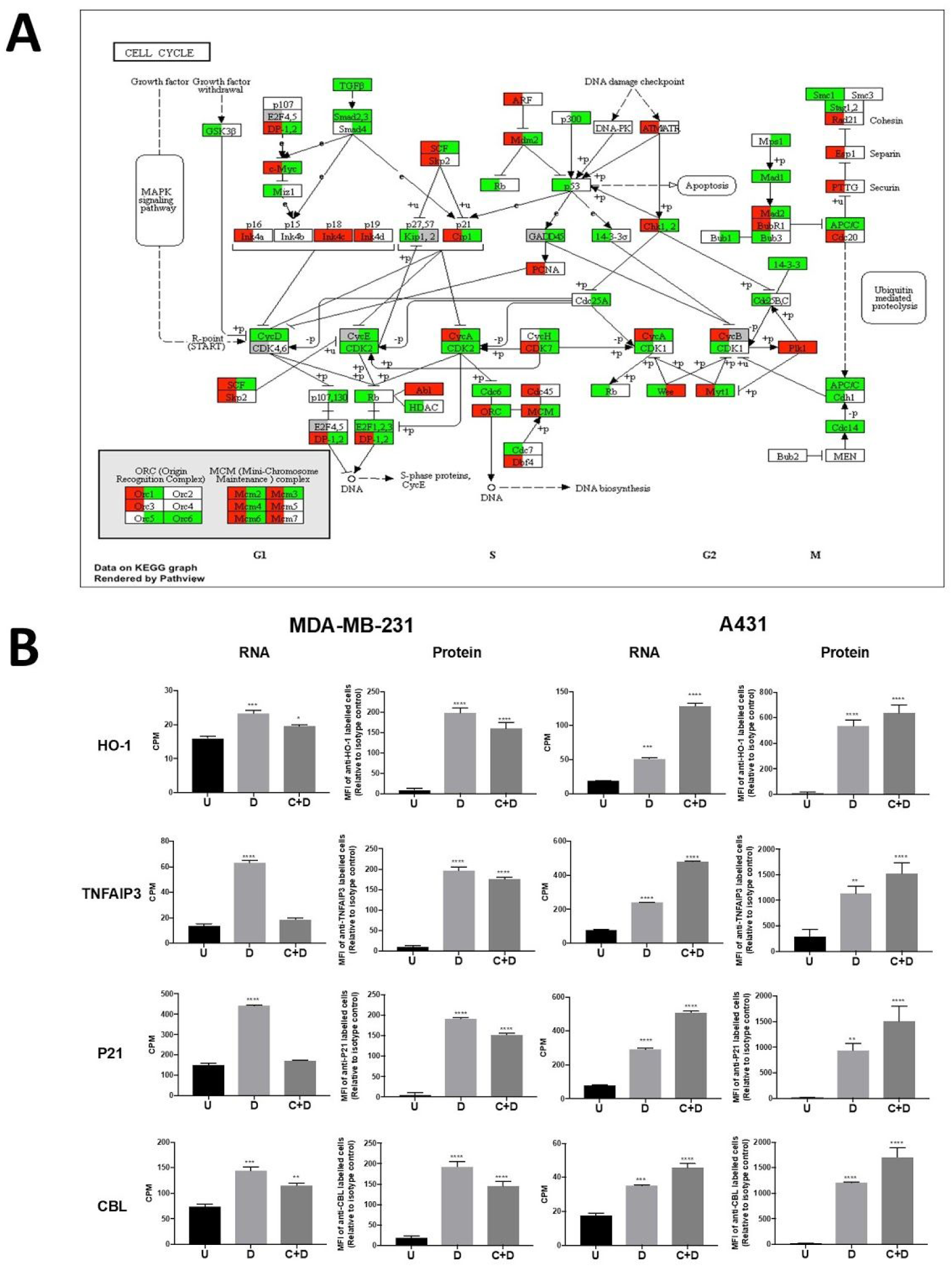
Differentially regulated genes shown in pathway and validation of selected gene regulation. A. Comparison of types of regulation for CKI with doxorubicin and 5-Fu in the “Cell Cycle” pathway. Left half of the rectangle for each gene represents CKI with doxorubicin in A431 cells and the right half represents CKI with 5-Fu in MDA-MB-231 cells. Red and green colors mean synergistic and antagonistic regulation, respectively. B. Validation of gene regulation at protein level. Four genes (HO-1, TNFAIP3, P21 and CBL) with opposite types of regulation in A431 and MDA-MB-231 cells identified by transcriptome sequencing were selected and validated by flow cytometry. ‘U’, ‘D’ and ‘U+D’ represent untreated, single chemotherapy drug treatment and CKI plus chemotherapy drug treatment, respectively. Data are represented as means ±SEM (n=9). Statistical analyses were performed between single drug treated or combined treated to untreated with one-way ANOVA (*p< 0.05, ***p < 0.001, **** p < 0.0001).

In order to validate the gene expression changes with different directions of regulation in the doxorubicin and 5-Fu treatment groups, we estimated protein abundance using flow cytometry for 4 proteins in group A. Overall, the protein level changes were consistent with gene expression levels from transcriptome analysis (Fig. 5B).

### Integrating information to select genes for validation

In order to select genes for experimental validation with bench experiments, we constructed the co-expression networks for genes in group A. 732 genes in 8 treatment groups were separated into 14 co-expression modules. By including data from the XTT and apoptosis assays, we calculated the correlation coefficients for each gene module with the phenotype results. The red, black and purple modules were more highly correlated with phenotype results than other modules. Because it had the highest correlation coefficients with both traits, genes in the red module (45 genes) were picked for further investigation (Fig. 6A).

**Figure 6:**
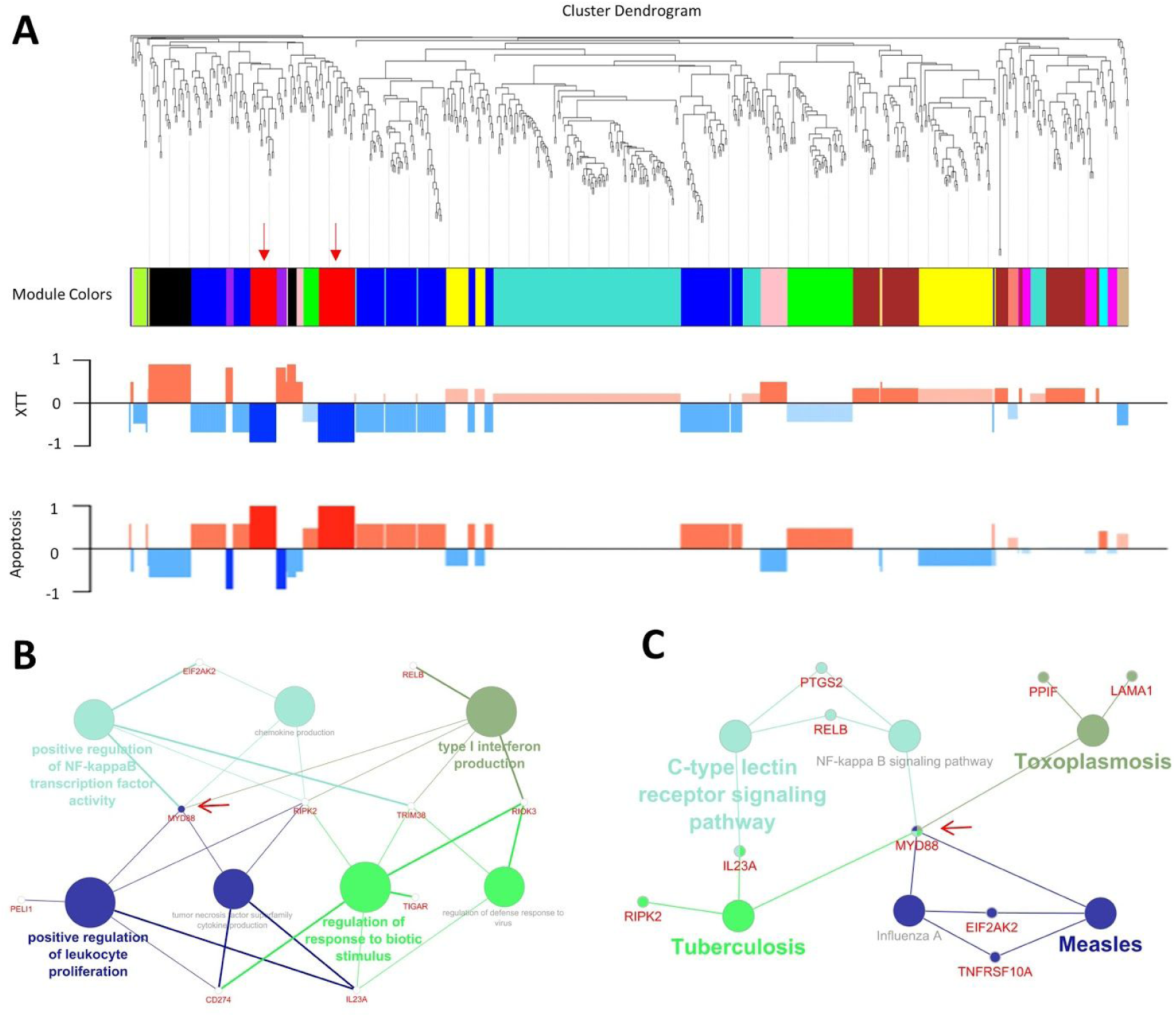
Co-expression analysis for genes in group A and related functional annotation. A. Clustering dendrogram for genes in group A and relationships with cell viability and apoptosis for each color module (red module is indicated by arrows). B. and C. Over-represented GO (Biological Process at 3rd level) and KEGG terms for genes in the red module.

Protein interaction, GO and KEGG analyses were performed for genes in the red module. From the protein interaction network, MYD88 was the most connected/interacting protein (Fig. 6B & C, Fig. 7A). Furthermore, it was commonly shared across different functions or pathways in the GO and KEGG analyses. Considering that MYD88 is upstream of NF-kappa B which itself regulates the cell cycle and other cancer related pathways, we selected it as a candidate for validation by inhibition.

**Figure 7:**
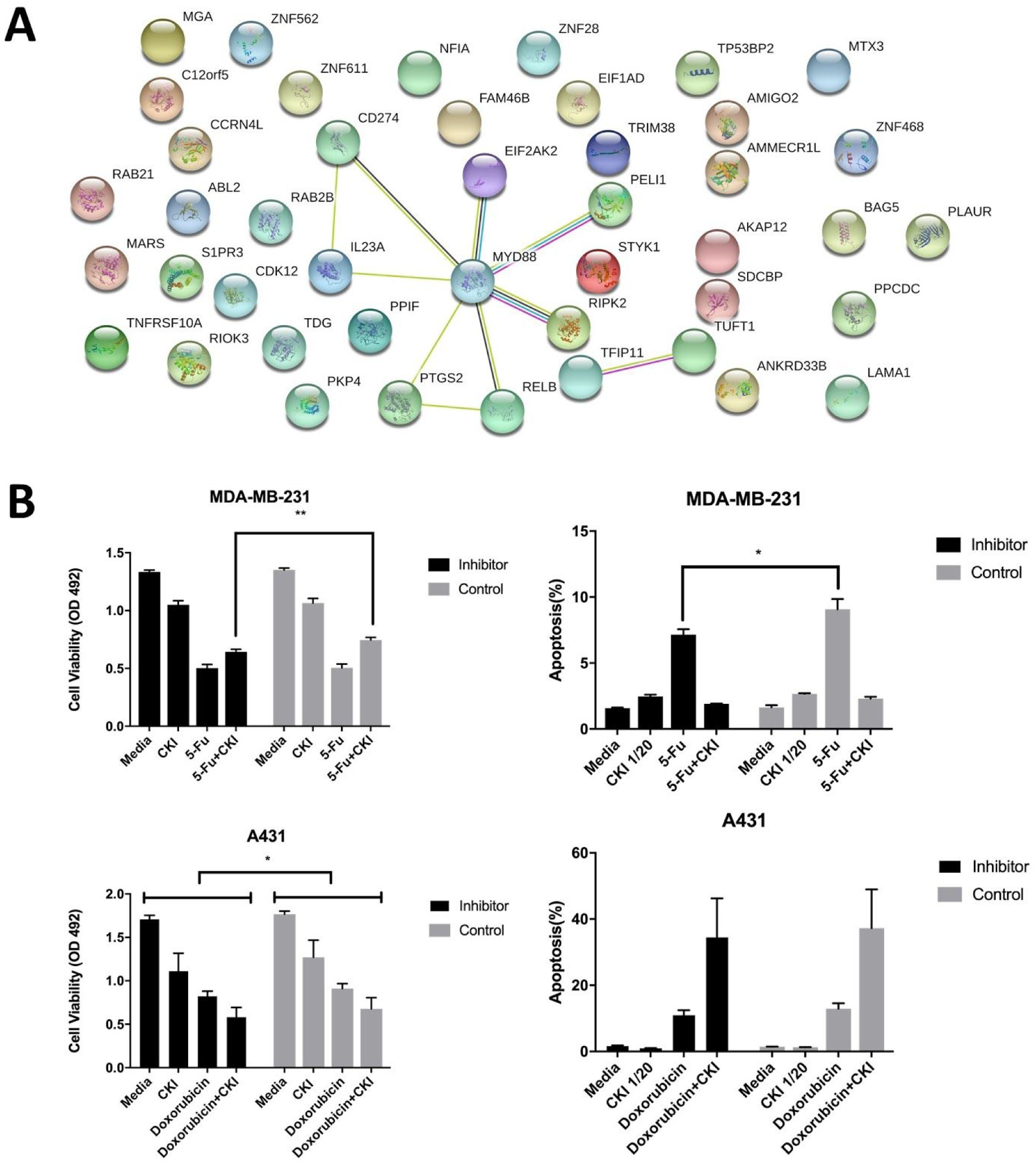
Validation of MYD88 function. A. All proteins for genes in the red module in Figure 6 and their interactions as based on the STRING database. B. Cell viability and percentage of apoptosis as a result of different treatments combined with MYD88 inhibitor peptide or control peptide. Results are represented as means ±SEM (n=9). Statistical analysis were performed with t-test (*p< 0.05, **p< 0.01).

### Inhibiting MYD88 partially affected the interaction between CKI and 5-Fu

To validate our analysis results, we performed the cell viability and apoptosis assays again and included an MYD88 inhibitory peptide or a control peptide (Fig. 7B). Because MYD88 is one of the key regulators in NF-kappa B pathway and occupies a central position in the red module, we expected that inhibiting it would reduce cell proliferation and the opposite effects from CKI on chemotherapy drugs. Results showed that for the MDA-MB-231 cells, inhibiting MYD88 does not affect the overall cell viability or apoptosis rates. However, compared to the control peptide, the inhibitor significantly reduced cell viability for 5-Fu and CKI combined treatment, by weakening the antagonistic effect of CKI. In the apoptosis assay, the apoptosis rate for 5-Fu treatment was significantly lower when treated with the inhibitor, also suggesting a similar reduction in the antagonistic effect. For the doxorubicin group, no significant changed interaction was found. However, unlike the MDA-MB-231 cell line, A431 cells were sensitive to the MYD88 inhibitor as shown by the overall lower cell viability values compared to the control peptide group.

In summary, we were able to dissect and characterise a DDI with transcriptome analysis. With CKI as a model, we identified candidate mechanisms behind its opposite effects compared to different chemotherapy agents and revealed potential interactions with them. We also identified and verified MYD88 as a target/key regulator for DDIs between CKI and anticancer drugs. These results demonstrate the value of our pipeline for characterising and understanding the molecular basis of DDIs.

## Discussion

Drug-drug interactions are one of the main reasons for adverse events associated with medication. The traditional pharmacokinetic methods for studying DDIs are inadequate for discovering potential side effects or explaining complicated interaction mechanisms. Furthermore, the complex components for many complementary medicines and herbal medicines that are often used in conjunction with pharmaceutical drugs pose a significant challenge research on DDIs. Although high-throughput omics-related techniques have been widely used for identifying novel disease biomarkers or potential drug targets ^16,17^, very limited research has applied them to the investigation of DDIs. Because transcriptome based approaches generate very large data sets, we adopted a hierarchical approach for our analysis of DDIs between CKI and chemotherapy drugs. First, instead of comparing every treatment sample to untreated control in order to identify DE genes, we decided to set the baseline for comparison of interactions as the main drug treatment. We then identified DE genes for the combined treatment based comparison to the main drug treatment. From the common set of DE genes found by comparing the main drug treatment to untreated control, and the combined treatment compared to the main drug treatment, we selected only genes that were differentially expressed and shared across the various comparisons. Second, we used “consistent directional regulation” to separate the DE genes between multiple treatments into positive and negative interaction classes. These are more informative with respect to drug-drug interaction than simply up or down regulation. This also allowed genes to be separated based on consistent directional regulation to focus the scope of investigation. Finally, we applied gene co-expression network analysis to provide useful information for candidate gene selection. These methods combined with typical gene annotation analysis and protein interaction analysis, provided a rich profile for investigation of DDIs.

Diseases treated with drug combinations are usually complex and related to multiple genetic pathways. Furthermore, multiple active compounds in drug combinations, such as herbal medicines can affect a variety of targets ^18 19^. Therefore, by integrating the effects of interacting genes, analysis of pathways or networks may provide more useful evidence for characterising the mechanisms of drug interactions. For the mechanisms of opposite effects generated by CKI combination with drugs, the pathways related to DNA synthesis and metabolism, like “Base excision”, “DNA replication” “homologous recombination”, are oppositely interacted between the two treatment groups. Related to both chemotherapy drugs down regulates genes in these pathways, the opposite interacted effects from CKI can enhance effects from doxorubicin while reduce 5-Fu’s effects. Closely linked to these pathways, “cell cycle” and “apoptosis” also have large differences in their manner of interaction between the two groups. By correlating with the results from cell viability and apoptosis assays, we can propose the opposite effects from CKI with doxorubicin or 5-Fu are primarily induced from pathways related to DNA synthesis and metabolism. As 5-fu and doxorubicin both target DNA replication and CKI’s cytotoxic effects have also been shown to increase DNA Double Strand Breaks [CITE JIAN’S BMC CANCER PAPER HERE] this indicates that therapeutic results from drug combinations targeting the same or similar bioprocesses can be very different compared to what we might predict ^13,20,21^.

Furthermore, by performing annotation analysis for functional or expression level clustering of gene groups, more information about interactions between CKI and chemotherapy drugs can be acquired. For group A, the annotation results are similar with opposite interacted results discussed in last paragraph, which is close related to phenotype results. In addition, the other three groups can help us to discover potential interactions which are not shown in our limited experiments. Group C displayed negative interaction of CKI to both doxorubicin and 5-fu and most annotation terms were belonged to organic biosynthetic and metabolic processes. Since many shared side effects from these two chemotherapy drugs are linked to disorders of metabolism, for example, cardiovascular and mucosal toxicity caused by cancer therapies are mainly caused by free radicals and oxidative stress ^22–24^. Therefore, although more validation is needed, the results for group C might support the clinical reports that CKI can reduce the adverse effects of chemotherapy and radiotherapy in cancer treatment. In addition, we observed two pathways “steroid biosynthesis” and “Fluid shear stress and atherosclerosis” that suggest that doxorubicin and 5-Fu affect atherosclerosis in a manner opposite to that of CKI. We could find no existing literature that would corroborate this finding. Our results indicate that transcriptome analysis can not only reveal candidate molecular mechanisms altered by specific drugs, but can also provide clues about potential drug-drug interactions.

Transcriptome analysis can provide a far more comprehensive and complex candidate gene list than traditional approaches used in drug-drug interaction research. This makes it difficult to screen target genes for further study because of the gene specific assays required. In group A, we generated a list of 732 genes, including heme oxygenase 1 (HO-1) and E3 ubiquitin-protein ligase (CBL), which are involved in metabolism pathways, that were oppositely regulated when CKI was combined with doxorubicin or 5-Fu. In addition, genes like tumor necrosis factor, alpha-induced protein 3 (TNFAIP3) and myeloid differentiation primary response protein (MYD88) from the NF-kappa B pathway and cyclin-dependent kinase inhibitor 1A (P21) in the cell cycle are regulated in the same manner. Although their functions are different, all these genes are important in carcinoma ^25–28^. By using gene co-expression and protein interactions analysis, we chose MYD88 as a proof of concept for validation as it was highly correlated with phenotype results and interacted with more proteins in its WGCNA color module than others. Our prediction was that inhibiting MYD88 would decrease the antagonistic effect of CKI on 5-Fu and this prediction was confirmed. By using our approach, transcriptome analysis can not only be used for generate comprehensive gene lists for candidate mechanisms, it can also identify specific, potential targets for modulating drug-drug interactions.

In summary, we introduced a pipeline to integrate omics techniques into research for DDIs. By using transcriptome analysis to identify candidate mechanisms that might account for CKI’s opposite effects on doxorubicin or 5-Fu in cancer cells, we have shown that our methods are effective and can be applied to complex situations, including drug interactions with complex mixtures or to compare different drug-drug interaction groups.

## Methods

### Cell culture and drugs

A431 and MDA-MB-231 cells were purchased from ATCC (VA, USA) and cultured in DMEM (Thermo Fisher Scientific, MA, USA) with 10 % fetal bovine serum (Thermo Fisher Scientific) at 37°C. with 5 % CO_2_. CKI (total alkaloid concentration of 26.5 mg/ml) was provided by Zhendong pharmaceutical Co.Ltd (China) and used at a final concentration of 1 mg/ml. Fluorouracil (5-Fu) and doxorubicin were purchased from Sigma-Aldrich (MO, USA) and used at final concentrations of 10 ug/ml and 1 ng/ml, respectively. MYD88 inhibitor and control peptides were synthesised by GenScript (Hong Kong, China) with the following amino acid sequences with purity > 95%^29,30^; inhibitor: DRQIKIWFQNRRMKWKKRDVLPGT and control peptide: DRQIKIWFQNRRMKWKK.

For all *in vitro* assays 6-well or 96-well plates were used. The seeding density for both A431 and MDA-MB-231 cells was 4 × 10^5^ m cells/well for 6-well plates. For 96-well plates, A431 cells were seeded at 8 × 10^4^ cells/well and MDA-MB-231 cells were 1.6 × 10^5^ cells/well. After seeding, cells were cultured overnight before being treated.

### Cell viability assay

ells were seeded in 96-well plates with 50 μl of medium. For the MYD88 validation assay, the inhibitor or control peptide was added at the same time as cell seeding. After overnight culturing, 50 μl of CKI and/or chemotherapeutic agent at appropriate concentration were added and incubated for 48 hours. In order to measure the cell viability, 50 μl of XTT:PMS (at 1 mg/ml and 1.25mM, respectively, and combined at 50:1 ratio, Sigma-Aldrich) was added and incubated 4 hours before detecting absorbance of each well with a Biotrack II microplate reader at 492 nm. Wells without cells were set up for each treatment for subtracting background absorbance.

### Cell cycle assay

Cells were cultured and treated in 6-well plates. After 48 hours of drug treatment, cells were harvested and stained with propidium iodide (PI) to examine cell cycle phases as previously described ^31^. Stained cells were acquired on BD LSRFortessa-X20 (BD Biosciences, NJ, USA) and the data were analysed using FlowJo software (TreeStar Inc., OR, USA).

### Flow cytometric quantification of protein expression

Cells were cultured in 6-well plates and treated with drugs for 48 hours. The cells were subsequently harvested and stained with antibodies to detect intranuclear/intracellular protein levels. The antibodies were purchased from Abcam (UK) unless otherwise indicated: rabbit anti-CBL and rabbit IgG isotype control (Cell Signaling Technologies) detected with anti-rabbit IgG-PE (Cell Signaling Technologies); mouse anti-p21 and mouse IgG2b isotype control detected with anti-mouse IgG-Alexa Fluor 488; rabbit anti-TNFAIP3-Alexa Fluor 488 and rabbit IgG isotype control-Alexa Fluor 488; rabbit anti-HO-1-Alexa Fluor 568 and rabbit IgG isotype control-Alexa Fluor 568. Data were acquired with a BD Accuri (BD Biosciences) and analysed with FlowJo software.

### RNA extraction and sequencing

After being treated with drugs in 6-well plates for 48 hours, cells were harvested and snap-frozen with liquid nitrogen then stored at −80 °C. Total RNA was isolated with the RNA extraction kit (Thermo Fisher Scientific) and quantity and quality were measured with a Bioanalyzer at the Cancer Genome Facility of the Australian Cancer Research Foundation (Australia) to ensure RINs > 7.0. Samples were sent to Novogene (China) and carried out on an Illumina HiSeq X platform with paired-end 150 bp reads.

### Transcriptome data analysis

Trim_galore (v0.3.7, Babraham Bioinformatics) was used to trim adaptors and low-quality sequences in raw reads with parameters: --stringency 5 --paired. Then trimmed reads were aligned to reference genome (hg19, UCSC) using STAR (v2.5.3a) with parameters: --outSAMstrandField intronMotif --outSAMattributes All --outFilterMismatchNmax 10 --seedSearchStartLmax 30^32^. Differentially expressed genes between two groups were calculated with edgeR (v3.22.3) and selected with false discovery rate (FDR) < 0.05^33^.

ClueGO was used to perform the GO and KEGG over-representation analyses with following parameters: right-sided hypergeometric test for enrichment analysis; p values were corrected for multiple testing according to the Benjamini-Hochberg method and biological process at 3rd level for GO terms34. Then Cytoscape v3.6.0 were used to visualise selected terms or pathways35. Regulation profiles for specific pathways were visualised with the R Pathview package36.

Co-expression network analysis was performed with WGCNA with “16” as soft thresholding power and “5” as minimum gene size for module reconstruction^37^. String (V11.0) was used to show protein interactions with 0.4 for minimum interaction score^38^.

