## Supplementary Figure Legends for "A new strategy for identifying mechanisms of drug-drug interaction using transcriptome analysis: Compound Kushen injection as a proof of principle"

Supplementary Figure 1: Cell viability of cancer cells treated with different drug combinations for 48 hours (MDA-MB-231 cells with doxorubicin and A431 cells with 5-Fu). Results are represented as means ±SEM (n=9). Statistical analysis was performed by comparing treatments to untreated (***p < 0.001, **** p < 0.0001)

Supplementary Figure 2: Comparison of types of regulation for CKI with doxorubicin and 5-Fu in the “Pathways in cancer” pathway. Left half of the rectangle for each gene represents CKI with doxorubicin in A431 cells and the right half represents CKI with 5-Fu in MDA-MB-231 cells. Red and green colors mean agonistic and antagonistic regulation, respectively.

Supplementary Figure 3: Multiple dimensional scaling (MDS) plot for MDA-MB-231 samples based on expression profiles of all genes (Untreated in black, CKI in red, 5-Fu in green and CKI+5-Fu in blue).

Supplementary Figure 4: Multiple dimensional scaling (MDS) plot for A431 samples based on expression profiles of all genes (Untreated in black, CKI in green, doxorubicin in blue, CKI+doxorubicin in cyan).

Supplementary Table 1: Mapping rates for each RNA-seq result.

Supplementary Table 2: DE gene lists for different comparisons.

Supplementary Table 3: Gene list for groups based on type of regulation (Group A-D) and their over-represented GO terms (count > 4 and P-value < 0.05).
