## Supplementary figures and images for "A new strategy for identifying mechanisms of drug-drug interaction using transcriptome analysis: Compound Kushen injection as a proof of principle"

### Supplemenetary Figure 1

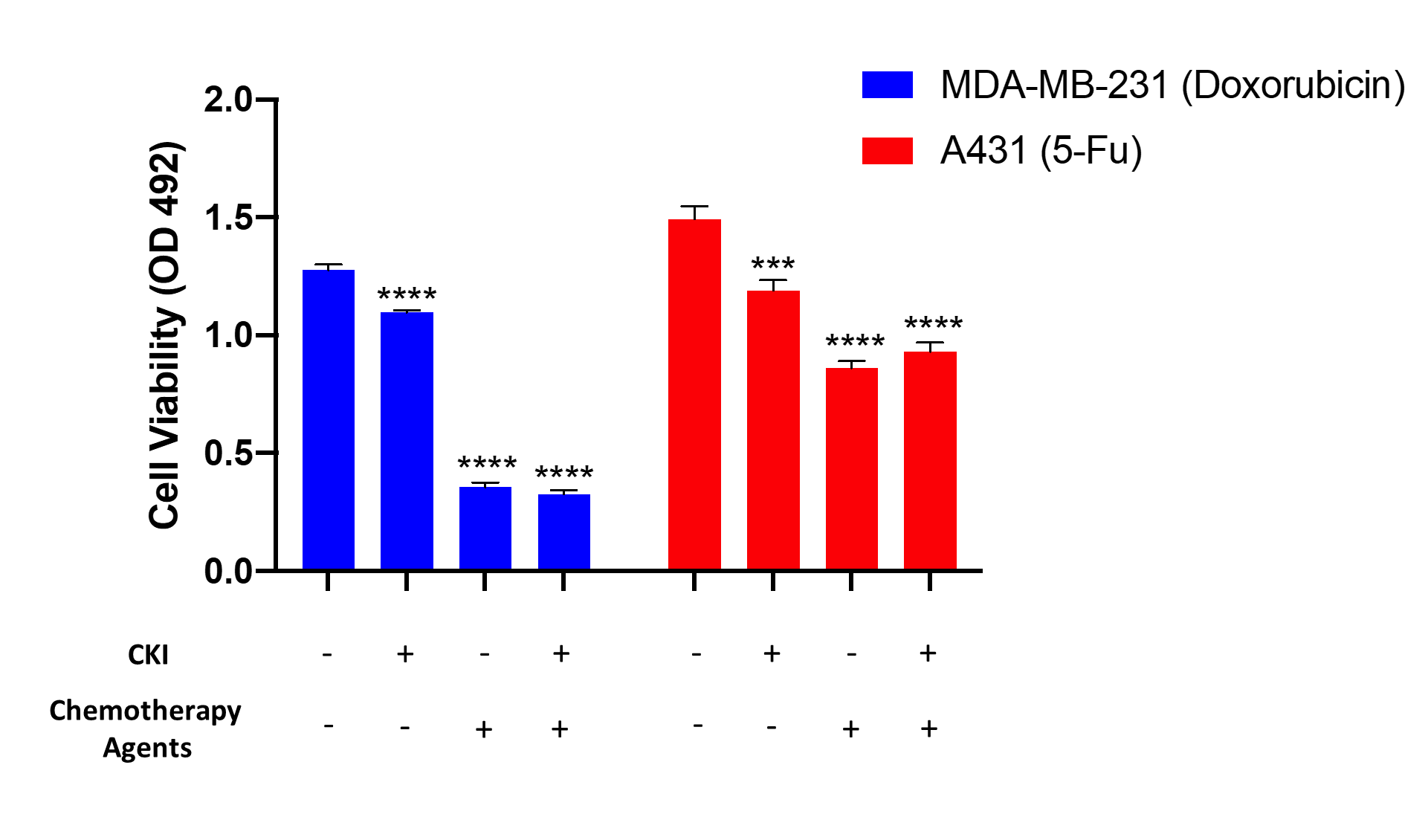

### Supplementary Figure 2

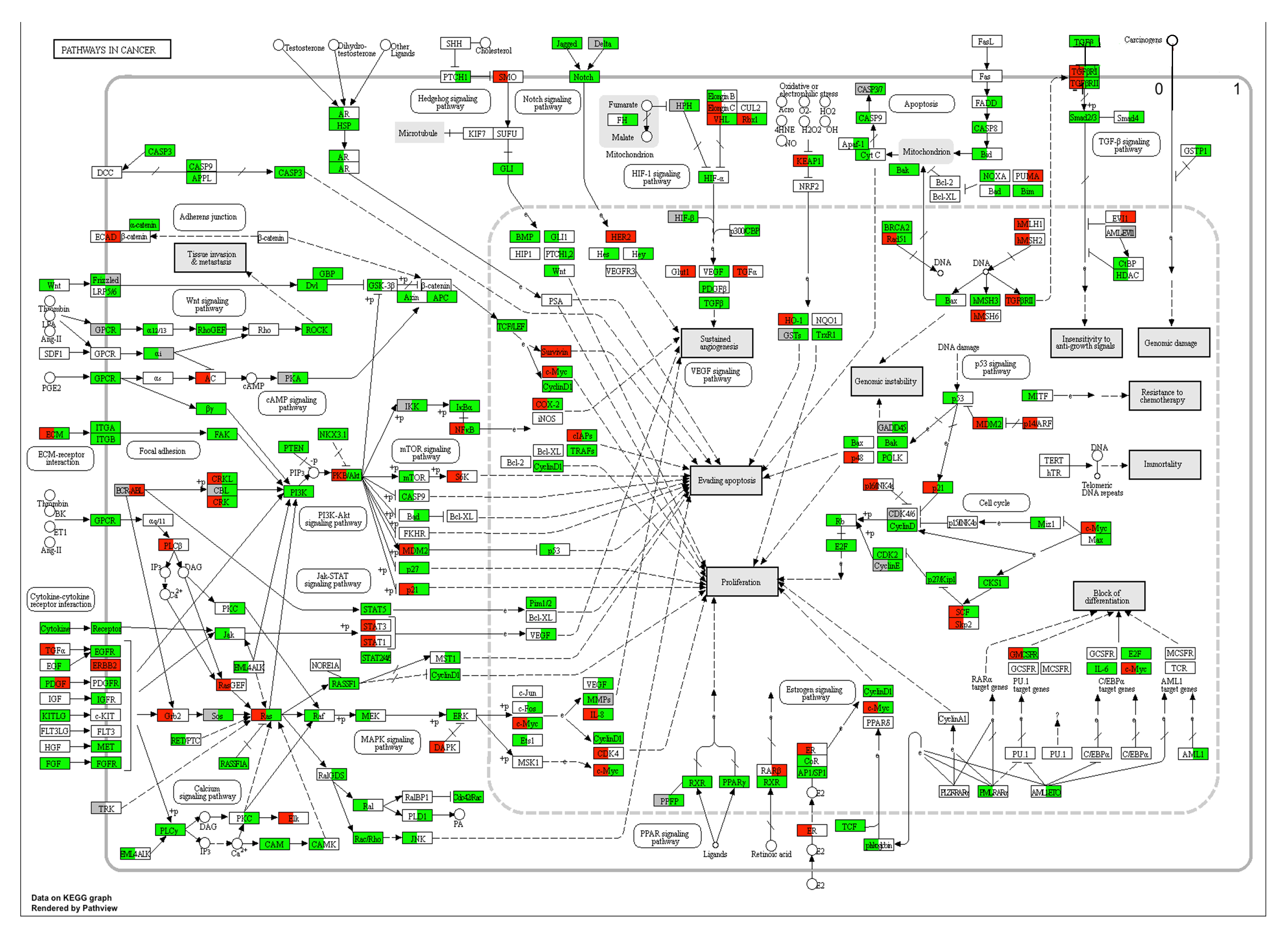

### Supplementary Figure 3

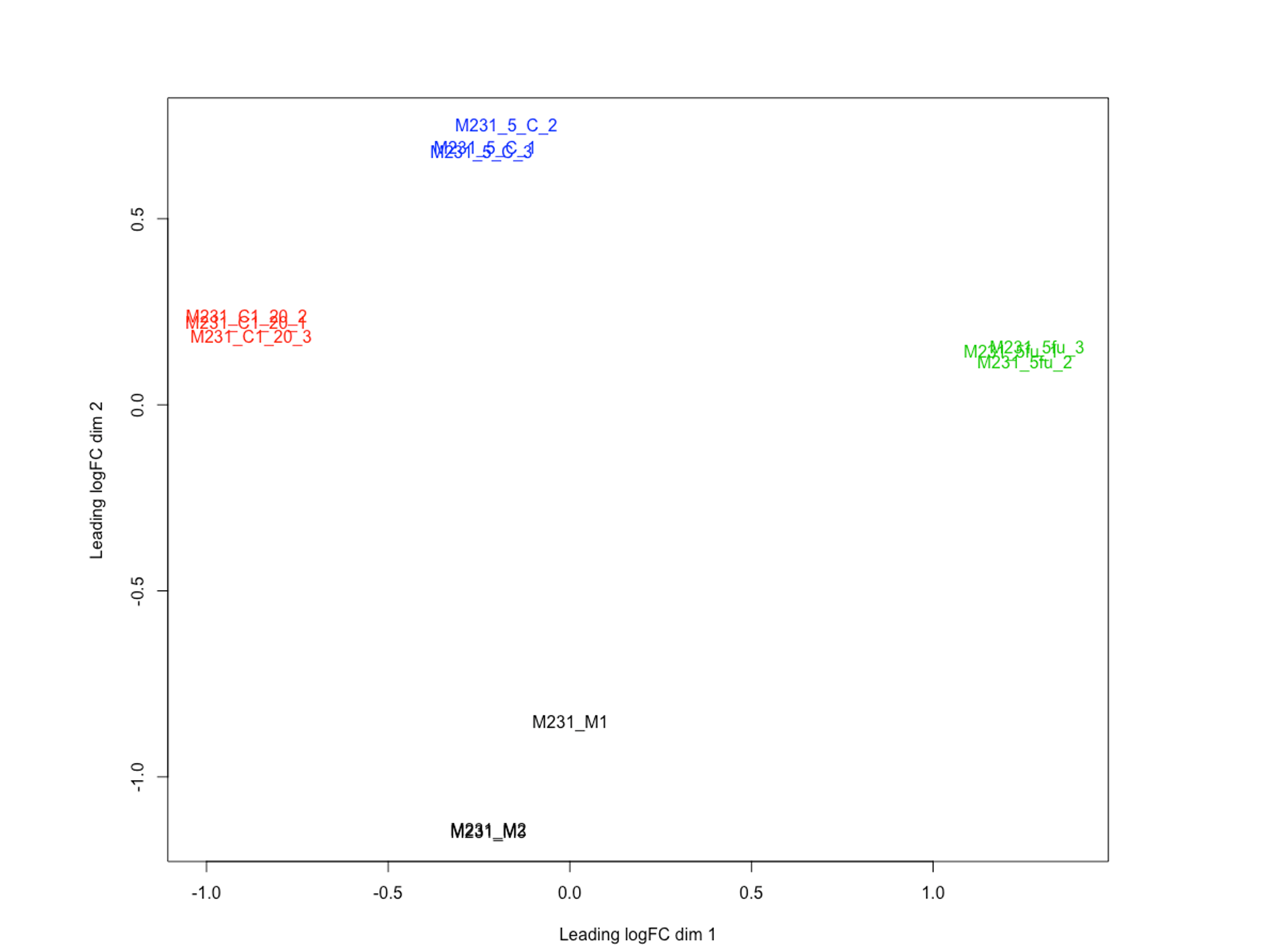

### Supplementary Figure 4

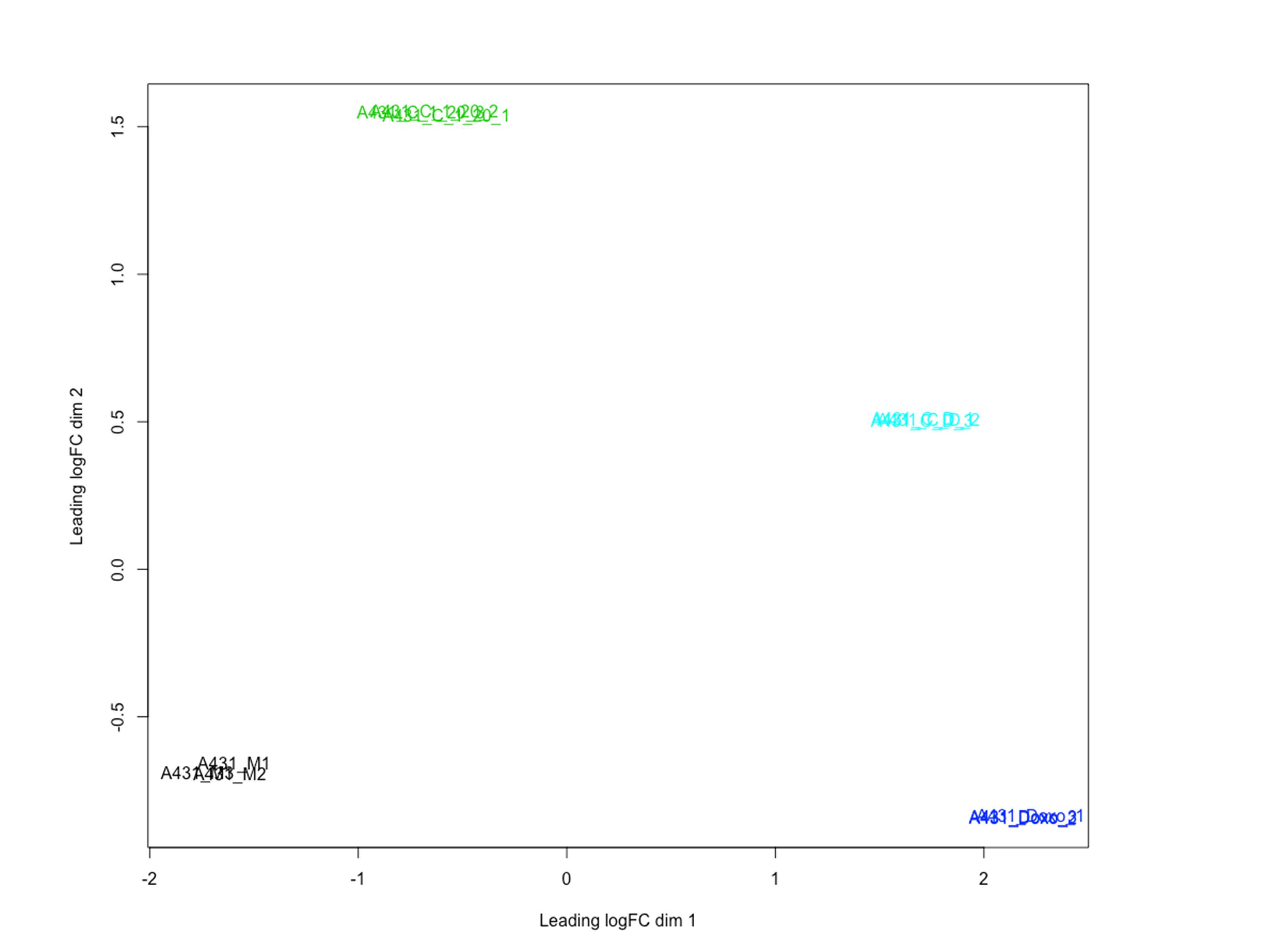
